# Serological and Molecular Investigation of Foot and Mouth Disease Virus and other animal pathogens at the Interface of Akagera National Park and Surrounding Cattle Farms between 2017 and 2020

**DOI:** 10.1101/2021.08.21.457188

**Authors:** Jean Claude Udahemuka, Gabriel Aboge, George Obiero, Ingabire Angelique, Natasha Beeton, Evodie Uwibambe, Phiyani Lebea

**Affiliations:** Centre for Biotechnology and Bioinformatics, University of Nairobi, P.O. Box 30197, Nairobi, Kenya; Department of Veterinary Medicine, University of Rwanda, P.O. Box 57, Nyagatare, Rwanda; Department of Public Health, Pharmacology and Toxicology, University of Nairobi, P.O. Box 29053, Nairobi, Kenya; Rwanda Agriculture and Animal Resources Board, P.O. Box 5016 Huye, Rwanda; TokaBio (Pty), Ltd, Pretoria, South Africa

**Keywords:** FMD, RT-LAMP, RT-PCR, Rwanda, Sequencing, SAT 2, Seroprevalence

## Abstract

**Background:** Foot-and-Mouth Disease Virus (FMDV) is a positive-sense RNA virus of the family of the picornaviridæ and responsible for the disease with the highest economic impact, the Foot-and-Mouth Disease (FMD). FMD is endemic in Rwanda but there are gaps in knowing the seroprevalence and molecular epidemiology. This study reports the FMD seroprevalence and molecular characterization of FMDV in Eastern Rwanda. Surveillance in FMDV wild reservoirs, the African buffaloes, was also carried out revealing the presence of other pathogens and commensals.

**Results:** The overall seroprevalence of FMD in the study area is at 9.36% in cattle and 2.65% in goats. We detected FMDV using molecular diagnostic tools such as RT-PCR and RT-LAMP and the phylogenetic analysis of the obtained sequences revealed the presence of serotype SAT 2, lineage II. Sequencing of oropharyngeal fluids collected from African buffaloes revealed the presence of several pathogens and commensals but no FMDV was detected in buffaloes.

The plethora of pathogens identified from the buffalo gut gives an idea of the health challenges faced by cattle keepers in Eastern Rwanda due to possible cross infectivity on wildlife-domestic animals interface regions.

**Conclusions:** We recommend further studies to focus on sampling more African buffaloes since the number sampled was statistically insignificant to conclusively exclude the presence or absence of FMDV in Eastern Rwanda buffaloes. The use of RT-PCR alongside RT-LAMP demonstrates that the latter can be adopted in endemic areas such as Rwanda to fill in the gaps in terms of molecular diagnostics. The identification of lineage II of SAT 2 in Rwanda for the first time shows that the pools as previously established are not static over time.

## Introduction

Foot-and-Mouth Disease Virus (FMDV) is a positive-sense RNA virus of the family of the picornaviridæ (1). Based on the most variable part of the capsid, the VP1, FMDV is classified into seven serotypes (SAT1, SAT2, SAT3, O, A, C and Asia1) which are also subdivided into topotypes (2,3). Vaccination against one serotype does not confer protection against a different serotype and multivalent vaccines are often used (4–6). Due to the constant change of this virus, a consistent molecular analysis is of paramount importance to improve the vaccines. Understanding the serological and molecular epidemiology of FMDV in Rwanda is very important because East Africa is considered to have the most difficult Foot-and-Mouth Disease (FMD) situation in the world (7).

In this study, we characterized the FMDV strains responsible for FMD outbreaks in Rwanda between 2017 and 2020 in cattle. Molecular diagnostics were performed using Reverse Transcription Polymerase Chain Reaction (RT-PCR) and the pen-side Reverse Transcription Loop-Mediated Isothermal Amplification (RT-LAMP) chargeable and portable machine, the Axxin T8 isothermal instrument. This was complemented by sequencing the strain responsible for the 2017 FMD outbreak. Moreover, we are presenting the serological situation of FMDV in large and small ruminants from the Eastern Province of Rwanda. These data will be crucial in policymaking and enforcement but also in studying risk factors. We also sampled wildlife, in African buffaloes (*Syncerus caffer*) known to be natural reservoirs of FMDV (8,9). Though we did not isolate FMDV in African buffaloes, we identified other pathogens and commensals. In Rwanda, there is no previously published research on FMD seroprevalence and the available FMD molecular characterization results are very old.

## Results

### Seroprevalence

We collected samples from adult animals as follows: 823 bovine sera samples and 188 caprine sera samples in 4 districts of the Eastern Province of Rwanda. The overall prevalence was 9.36% (77/823, CI_95%_ : 98.0-99.9) in cattle and 2.65% (5/188, CI_95%_ : 96.9-99.9). The seroprevalence distribution in bovine was as follows; 8.6% (55/639) in Bugesera and 11.96% (22/184) in Nyagatare. In caprine, the seroprevalence was at 2.77% (4/144) in Bugesera and 2.27% (1/44) in Kayonza.

### Reverse Transcription Polymerase Chain Reaction (RT-PCR)

The 2017 FMD outbreak samples were analysed by RT-PCR. The 1-step RT-PCR assays performed on the samples collected from the field samples demonstrated 6 out of 9 oropharyngeal (OP) samples positively identifying infection with the FMD virus. Select PCR positive samples are displayed in figure 1 and an increase in fluorescence intensity (Relative Fluorescence Units or RFU) was detected before the threshold of 32.0 cycles of amplification. Sample 8 and 26 representing Gatsibo and Nyagatare districts respectively were particularly chosen for further analysis. Figure 2 illustrates the triplicates results of select samples.

**Figure 1:**
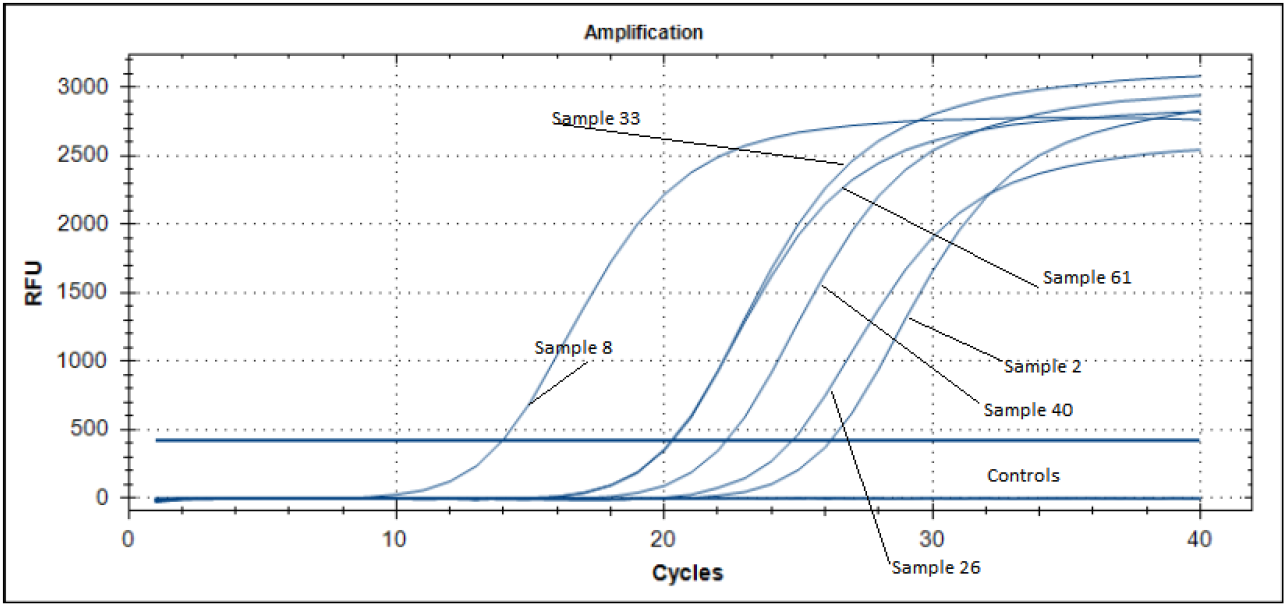
One-Step RT-PCR analysis. Amplification curves illustrating some of the select positive samples from different animals in both the Gatsibo and Nyagatare regions.

**Figure 2:**
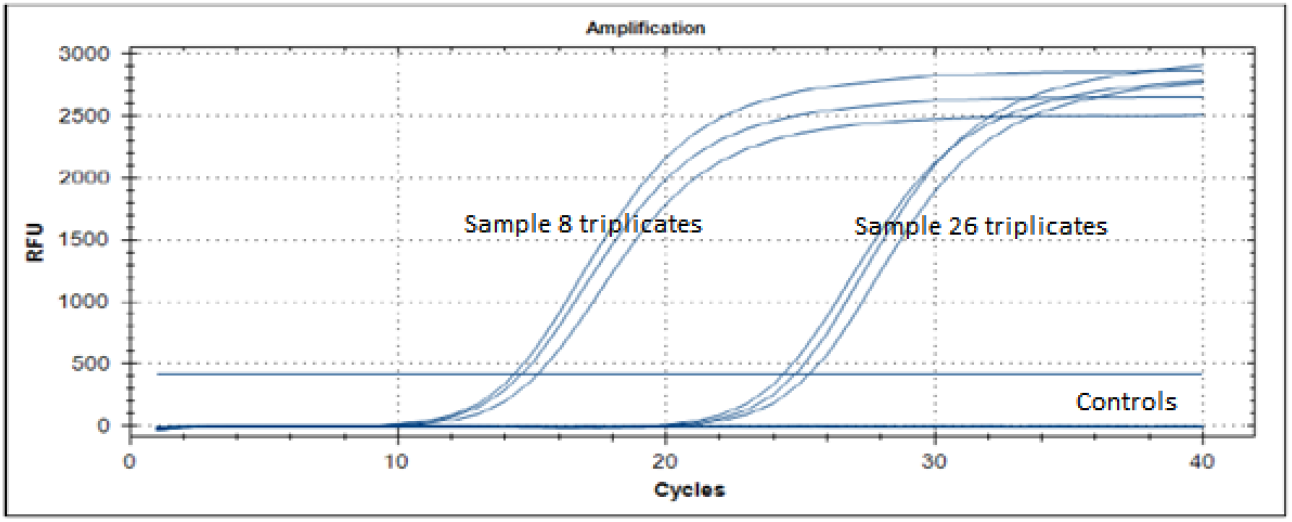
Amplification curves illustrating the detection of FMD viral RNA in the reaction, using OIE recommended TaqMan probes. The early triplicates were from sample 8 (Gatsibo) while the later threshold triplicates were from sample 26 (Nyagatare). The non-template controls are illustrated by a straight line around the zero FU level.

During the 2020 FMD outbreak, we collected samples from Kayonza district. Out of 19 blood samples, six (samples 7, 8, 9, 15, 16 and 18) showed bands (figure 3) for the gel-based PCR analyses using SAT 2 specific primers (Table 2) with sample number 6 having a faint band. Other sets of primers for serotypes SAT1, O and A did not reveal the presence of FMDV in those samples.

**Figure 3:**
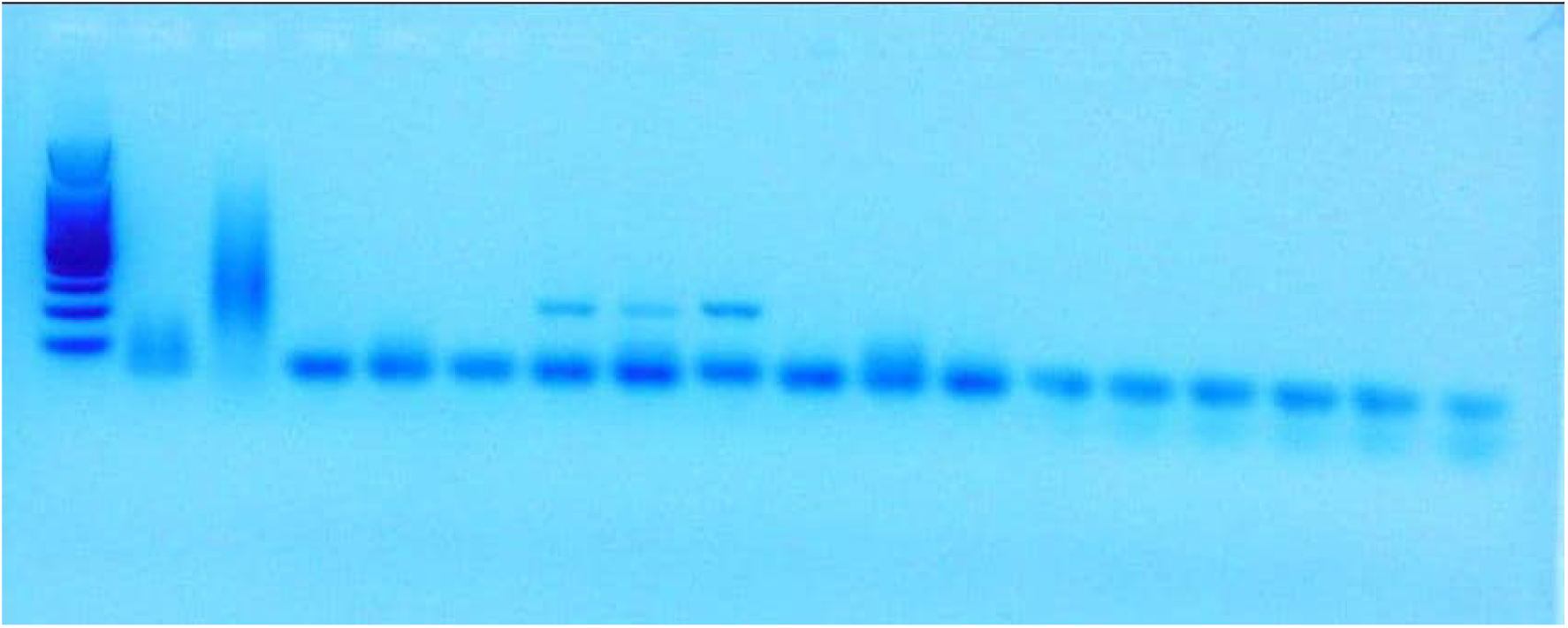
Electrophoresis results for the 2020 FMD outbreak in Kayonza district showing positive samples to FMDV SAT 2.

### Real-time reverse transcription loop-mediated isothermal amplification (rRT-LAMP)

The RT-LAMP assay analysis revealed positive FMD detection consistent with results revealed by real-time PCR profile detection from the samples collected in the field. The time trial fluorescence graphs are represented in figure 4 and the gel-based detection in figure 5.

**Figure 4:**
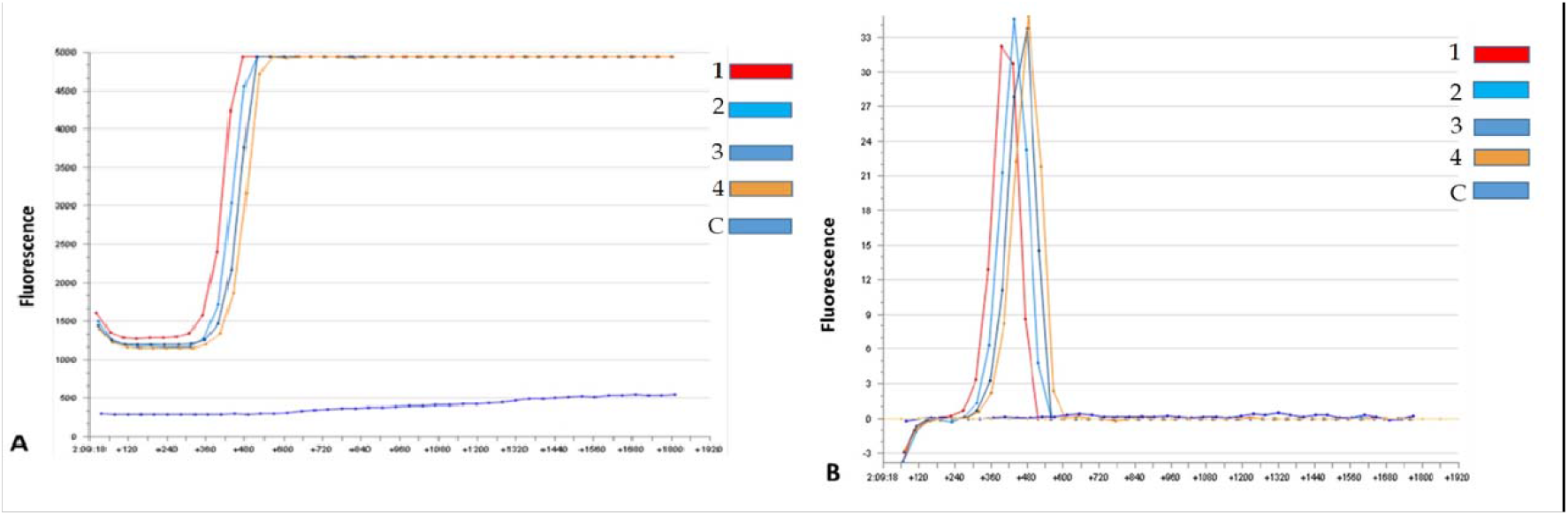
RT-LAMP results of field samples. Fig. 4.a describes the time trial of fluorescence detection of field samples and fig. 4.b shows the second derivative graphs of the fluorescence of the same samples collected in Eastern Rwanda during an outbreak.

**Figure 5:**
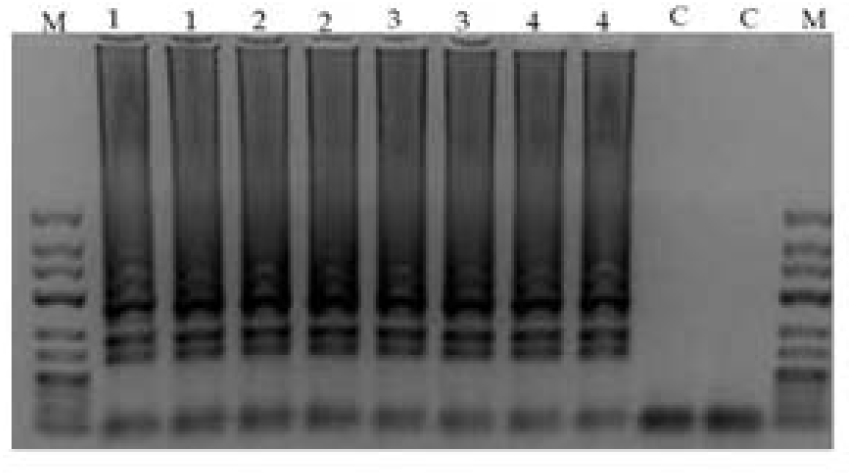
Agarose gel detection of RT-LAMP amplicons of selected PCR FMD positive duplicate samples from Nyagatare and Gatsibo.

### Sequencing

From the oropharyngeal fluids (OPF) sampled from A. buffaloes, we did not observe the presence of FMDV. However, we detected other pathogens and commensals (Figure 6), suggesting that they may be reservoirs of other infectious diseases of interest that can infect domestic ungulates in Rwanda.

**Figure 6:**
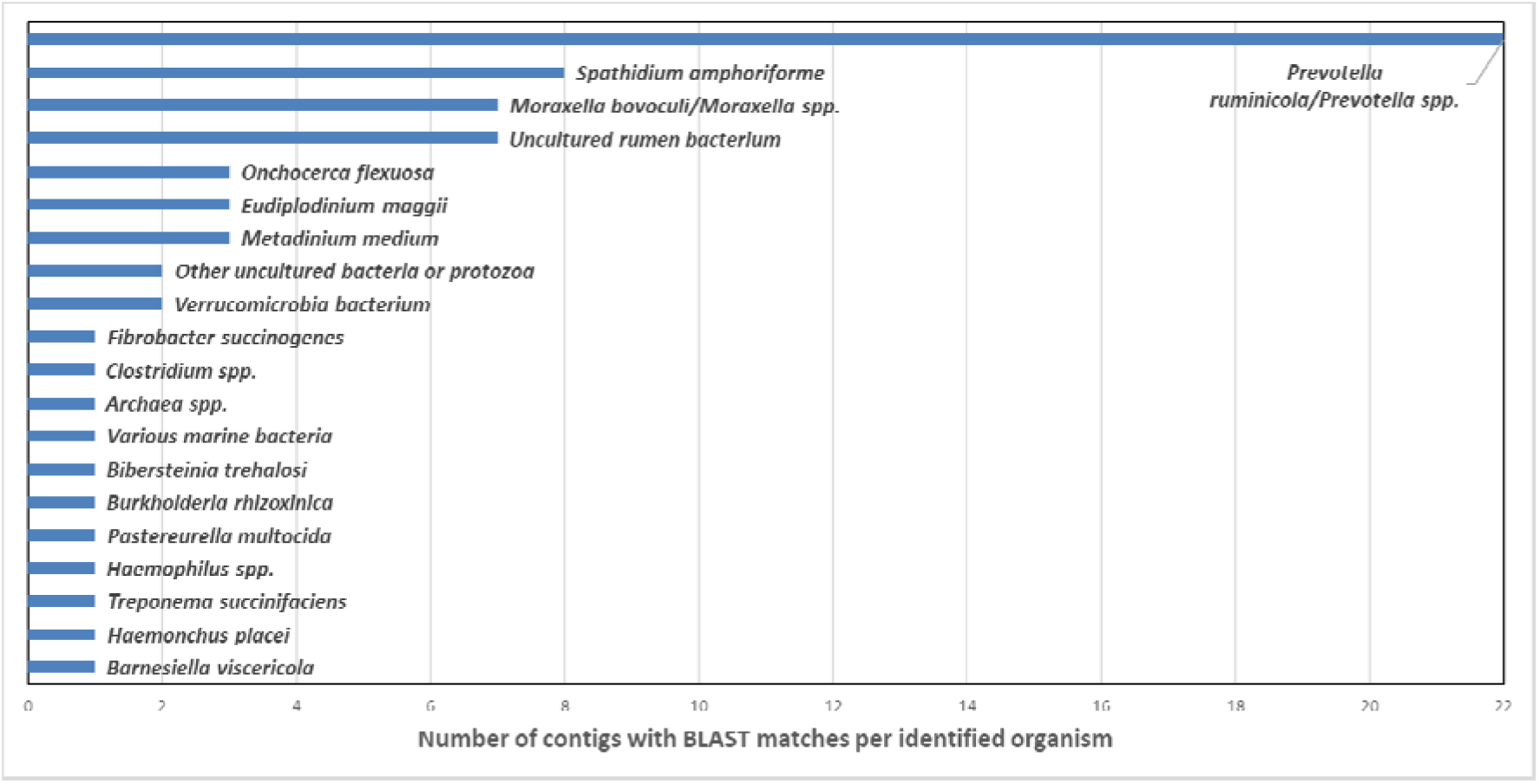
Bar chart showing the number of sequenced contigs across all seven African buffaloes with BLAST matches to species listed in the NCBI database (https://blast.ncbi.nlm.nih.gov/Blast.cgi).

FMDV whole genome was sequenced from OPF clinical samples collected from cattle and the sequences are available upon request to the corresponding author. Phylogenetic analyses of VP1 proteins from this study and prototypes available online revealed that topotype II of SAT 2 was the causing virus of the 2017 FMD outbreak in Rwanda. This study’s VP1 sequences were very similar to prototypes isolates SAT2/ZIM/7/83* (AF136607) and SAT2/ZIM/5/81 (EF134951) belonging to topotype II of serotype SAT 2 (figure 7). The prototype isolates previously identified in Rwanda (AF367134) and former Zaire (DQ009737) representing the topotype VII are not close to this study’s isolates.

**Figure 7:**
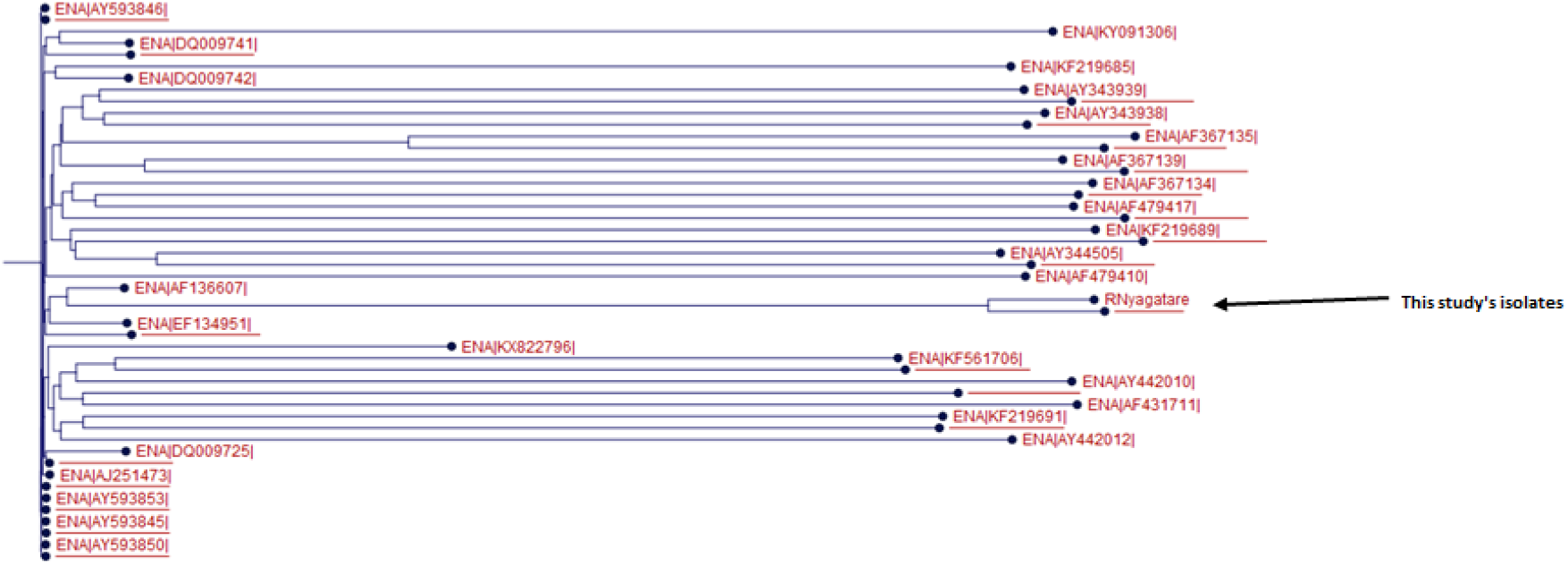
A phylogenetic tree of FMDV VP1 from Rwanda’s 2017 outbreak and SAT 2 prototypes

Pairwise comparison between isolates SAT2/ZIM/5/81 and SAT2/ZIM/7/82* and this study’s isolates, as illustrated in table 1, are all beyond the threshold of 80% (3).

**Table 1:** Pairwise comparison between FMDV VP1 from Rwanda’s 2017 outbreak and SAT 2 lineage II prototypes

|  |  | 1 | 2 | 3 | 4 |
| --- | --- | --- | --- | --- | --- |
| Isolate 1 (RGatsibo) | 1 |  | 0 | 0 | 2 |
| Isolate 2 (R Nyagatare) | 2 | 5 |  | 0 | 2 |
| EF134951 | 3 | 89 | 90 |  | 2 |
| AF136607 | 4 | 92 | 92 | 89 |  |

**Table 2:**
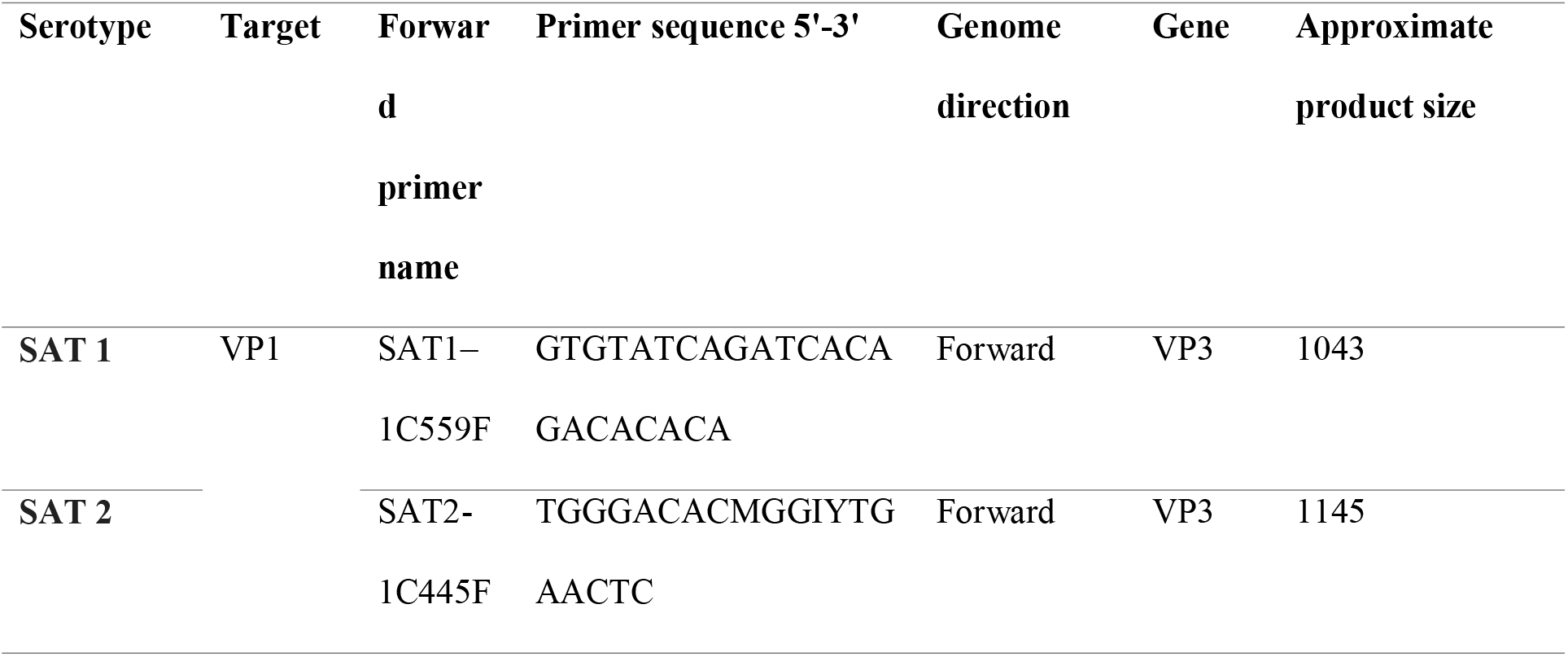
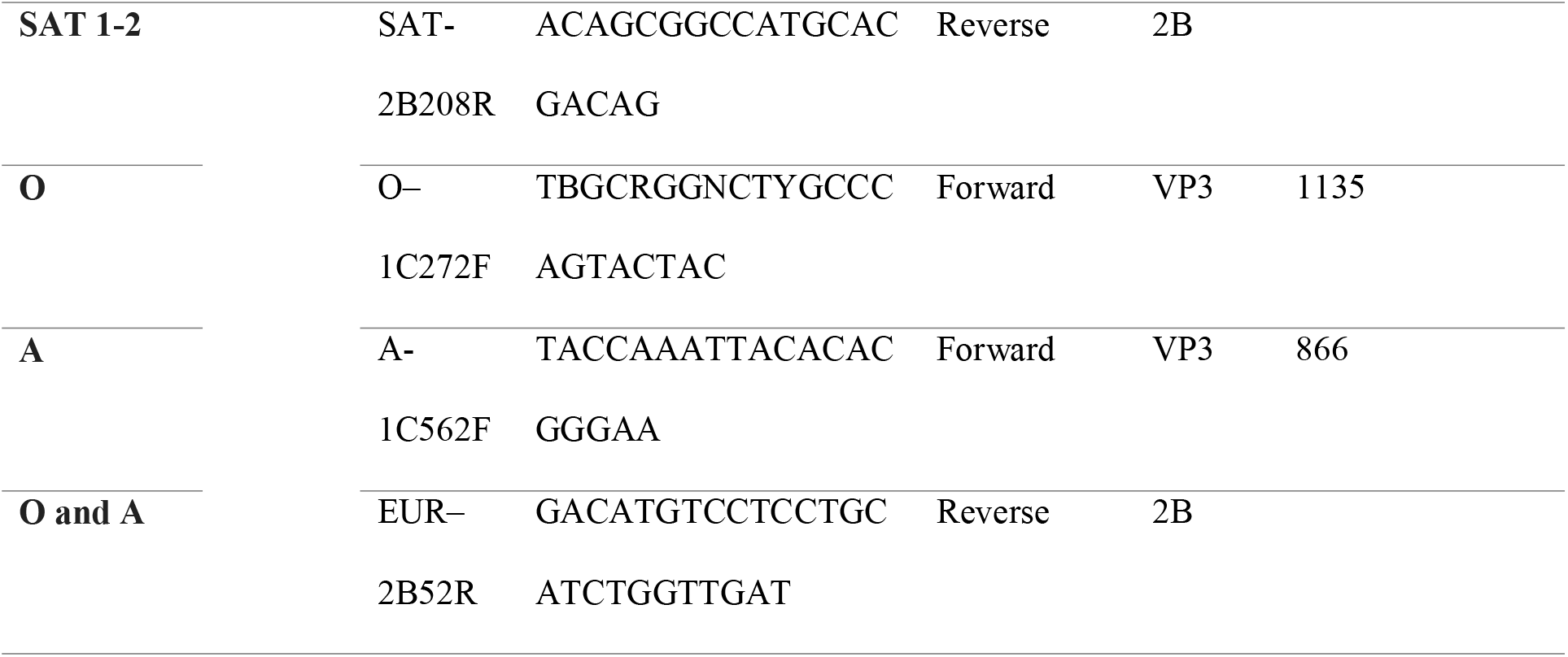
List of oligonucleotide primers used for a reverse-transcription polymerase chain reaction in this study (49)

## Discussion

### Seroprevalence

Current vaccines against FMDV are purified from non-structural proteins (NSP), this means that if

NSPs are detected in sera samples they are exclusively results of prior infection and not vaccination (10). The detection of antibodies to NSP such as the highly conserved 3ABC is widely used to determine the FMDV seroprevalence (11). Recently, we were not able to get information on seroprevalence for the study on FMD risk factors in Eastern Rwanda (12), therefore this study sets a baseline to get first-hand FMD seroepidemiological information in cattle and small ruminants in Eastern Rwanda. With 3ABC ELISA (Enzyme-Linked Immunosorbent Assay), antigens against FMDV were detected at the prevalence of 9.36% and 2.65% in cattle and in goats respectively. The higher prevalence in cattle may be explained by the fact that viral replication is expected to be more luxuriant and more persisting in cattle than in small ruminants (13). The higher prevalence of FMDV in bovine serum from Nyagatare, compared to Bugesera samples, may be due to the closeness of the Nyagatare district to wild FMD-susceptible animals. Therefore, a sero-surveillance study in wild FMD-susceptible animals would be valuable. Moreover, we observed that visited farms in Bugesera have more on-site resources such as water and fodder decreasing the risk of animals roaming outside the farm; this also may contribute to lower seroprevalence in Bugesera compared to Nyagatare.

### African buffalo samples

We did not detect FMDV in the seven samples of the African buffaloes randomly selected from the Nyamirama area inside the Akagera National Park in Rwanda. In Southern African countries, a pattern of cross-contamination has been established between livestock and wildlife animals considered as natural reservoirs, particularly African buffaloes (*Syncerus caffer*) (14,15). In that region, physical separation consisting of electro-fencing of parks and movement restriction combined with effective vaccination campaigns (16,17) have assisted in reducing outbreaks from cross-contamination and FMD control. However, in East Africa, few studies have been carried out on the circulating strains of FMDV, which constitutes a knowledge gap that prevents any conclusions that such measures have been similarly effective in this region. In 1979, SAT 3 was detected in African buffaloes in Queen Elizabeth National Park and in a healthy long-horned calf that grazed near the park.

Several reports have demonstrated that African buffaloes play an important role in the epidemiology of the SAT serotypes of FMDV, although only SAT3 has been identified thus far in buffaloes (18). Recent studies have shown that there are likely various routes of FMDV transmission from wildlife to livestock and vice-versa (19–21). However, among the few studies that have focused on FMD in African buffaloes, some have proposed that some outbreaks may be of a different origin, such as unregulated cattle movement as well as the porous borders in the region (19). In the present study, the sampled buffaloes had been enclosed inside the park and separated from farm access with an electrical fence since 2013 (22). Although there is a need for a greater number of buffalo samples to be analysed, the results of our study suggest that the series of outbreaks observed in Eastern Rwanda between 2000 and 2017 may be arising from transboundary and intra-national livestock movements as well as proximity with unvaccinated small ruminants. These factors would contribute to virus persistence in the area because FMDV can be recovered in small ruminants up to 9 months post-infection (23).

### Cattle samples

Regions in Eastern Rwanda, at the border between Uganda and Tanzania, are at risk of transnational circulation of the FMD virus. We typed the strain responsible for the 2017 and 2020 FMD outbreaks as SAT 2, this serotype is the most predominant in Sub-Saharan Africa (24,25). Considering the samples from Rwanda that have been typed previously, SAT 2 seems to be the most predominant serotype in the last 20 years in Rwanda as well (12). However, in the region, it comes second alongside serotype A and after serotype O (7,26–29). Across the border, in Western Tanzania (Kyerwa and Karagwe districts) and Western Uganda (Isingiro district), SAT 2 has been responsible for several FMD outbreaks (30–32). We hypothesize that if these two regions with Eastern Rwanda that make a triangle with transboundary animal movements are not controlled, they can contribute to the circulation of several strains of FMDV and other similar transboundary animal disease pathogens.

### Reverse Transcription Polymerase Chain Reaction (RT-PCR)

Amplification using specific primers targeting FMDV SAT2 serotype revealed the presence of serotype SAT 2 in the OP samples. In Kenya, most of the outbreaks have been caused by serotypes O and SAT2 (33). Among the SATs serotypes, SAT2 is the most predominant in East Africa in general (30) and in Rwanda in particular where the last serotype O outbreak was reported in 2004(34). We hereby report the shreds of evidence that SAT2 was responsible for the 2017 and 2020 FMD outbreaks. Therefore, special consideration should be given to vaccines containing SAT 2 strains isolated in the region if a regionally coordinated vaccination campaign is to be carried out.

### Real-time reverse transcription loop-mediated isothermal amplification (rRT-LAMP)

This method relies on auto-cycling strand displacement DNA synthesis that is performed by DNA polymerase with high strand displacement activity and a set of two specially designed inner and two outer primers (35).

FMDV constant molecular diagnostics allow a better understanding of circulating strains for a smarter vaccination. In this study, we have confirmed what other studies have established that LAMP technology can be an alternative to traditional PCR. Our LAMP results have shown that SAT 2 serotype was responsible for the 2017 outbreak, this in concordance with PCR results. This assay has been previously used in Southern Africa successfully on FMDV SATs serotypes (36).

The LAMP technology can be very useful in point-of-care detection during an active outbreak to detect FMDV while sampling is still going on. The Axxin T8 being a portable device with batteries making it suitable to be used in mobile laboratories and since RT-LAMP skips the cDNA synthesis step and the extraction step for some samples such as blood and cell cultures (37,38), it is much more fit for Low-and-Middle Income Countries and decreases the risk of contamination (39). RT-LAMP has proved to detect positive samples and within 30 minutes we were able to molecularly diagnose FMDV which would take a much longer time with laborious manipulations when diagnosing with RT-PCR. Also, RT-LAMP is better for blood samples because it is not hindered by inhibitory substances (39). Recently, Mahapatra *et al.* have demonstrated that LAMP’s concordance with RT-qPCR reaches 100% (40). This pen-side, rapid and cost-effective technology presents an alternative molecular diagnostic to RT-PCR and can be of great importance during outbreaks. The democratisation of such technologies would fill the gap of lack of proper knowledge on circulating strains in Eastern Rwanda.

### Bioinformatics analyses

The presence of SAT 2 lineage VIII has been previously identified in Rwanda (isolate SAT2/RWA/1/00, accession number AF367134) and in the neighbouring Democratic Republic of the Congo (isolate SAT2/ZAI/1/74, accession number DQ009737) (41,42). However, this study revealed the presence of lineage II, traditionally localized in Southern Africa and close to isolates SAT2/ZIM/7/83* (AF136607) and SAT2/ZIM/5/81 (EF134951). Recently, the introduction of new lineages in pools where they were not previously identified has occurred (43) due to the increase of transnational trade and traffic of people and animals. This can explain our results of finding lineage II of SAT 2 in East Africa. With the rise in African free trade and globalisation, specialists of animal disease control should be more cautious expecting more outbreaks of diseases or strains in areas they were not traditionally localised.

## Conclusion

While there were no FMDV pathogens isolated in African buffaloes, the whole genome sequencing revealed the presence of other pathogens that can cross infect cattle. The plethora of pathogens identified from the buffalo gut gives an idea of the health challenges faced by cattle keepers in Eastern Rwanda due to possible cross infectivity on wildlife-domestic animals interface regions. We recommend further studies to focus on sampling more African buffaloes since the number sampled was statistically insignificant to conclusively exclude the presence or absence of FMDV in Eastern Rwanda buffaloes.

The 2017 and the 2020 FMD outbreaks in Eastern Rwanda were caused by SAT 2 serotypes and the VP1 analysis of the 2017 sequences the presence of SAT 2 lineage II for the very first time in Rwanda, this highlights the probable role played by transboundary animal movements. In this case, one could not say whether it is of wildlife origin or domesticated animal trading that occurs from time to time.

It is noteworthy to recognise that RT-LAMP was used in the endemic areas of Eastern Rwanda as a rapid, and cost-effective alternative method for field detection. Additionally, this technology usage did not require high molecular biology skills to operate. This ability to run the RT-LAMP in the field made the technology suitable to be manipulated by a bigger number of animal health workers. We had previously successfully tested the RT-LAMP technology against other FMD serotypes, but testing this technology against SAT 2 is more meaningful for East Africa.

## Materials and Methods

### Source of samples and sampling techniques

We randomly immobilized seven mature African buffaloes (*Syncerus caffer*) in the Nyamirama area inside the park. We tranquilised buffaloes from a 4x4 vehicle or on foot with a 2 mL Pneu-Dart and a 3.8 cm barbed needle using a spring-loaded pole syringe (Dan-Inject) JM Special dart gun with a 13 mm barrel. We darted all animals in the hindquarters with 8 mg etorphine and 48 mg azaperone (44) and they all went down in sternal recumbence. We covered the face plugged the ears with a tissue, to avoid early wake-up. We injected 150 mg of Ketamine to increase muscle relaxation and jaw movements. We added a combination of an additional 200 mg ketamine IV and 2 mg etorphine with 40 mg azaperone IM to early woken up animals. After sample collection, buffaloes were roused with up to 20 mg diprenorphine and 100 mg naltrexone given intravenously (44,45) and all recovered well. Throughout the sample collection, we monitored the buffaloes’ breathing and no animal needed a respirator stimulant or partial antagonism. We collected OPF and scraps using a probang cup and transferred the samples to sterile tubes containing an equal amount of transport media. We inserted a probang cup in the OP tract and vigorously passed it with back-and-forth movements at least 5-10 times between the first portion of the oesophagus and the back of the pharynx. Tubes had equal amounts of glycerol and 0.04 M phosphate buffer (pH 7.2–7.6) containing 1× antibiotic–antifungal mixture (Thermo-Fisher Scientific, Johannesburg, South Africa) (46,47). Between OPF collection from one animal to the next using the OIE three-bucket system slightly modified (46,47). We washed the probang cup in a bucket containing 0.3% citric acid, rinsed it in another bucket with water and lastly disinfected the cup in Phosphate-buffered Saline. The sample tubes were topped up to contain an equivalent volume of transport medium to that of the sample.

Following the described methodology above, in July 2017, we collected from crossbred (Ankole x Jersey) cattle 9 OPF and scraps from cattle in the Gatsibo district during the 2017 FMD outbreak and 53 blood samples collected in EDTA tubes in Kayonza district during the FMD outbreak in May 2020. We transported samples in cooler boxes on ice from the field to the Virology Laboratory of the Rwanda Agriculture and Animal Resources Development Board located in Kigali, Rwanda and stored them at −80°C until further processing.

Alongside the sequences produced by this study, we retrieved sequences from the National Center for Biotechnology Information (NCBI). We targeted sequences identified as SAT 2 from samples collected in neighbouring countries namely, Uganda and Tanzania. We provided details of the retrieved sequences in annexe I.

### Serological analyses

Sera samples were randomly collected from three districts of the Eastern province during surveillance. Samples were stored at −20°C for less than one week before analysis. We used the ID Screen® FMD NSP Competition Kit (ID.Vet, Grabels, France) according to the manufacturer’s instructions to detect the non-structural protein 3ABC in serum. The test was applied to samples from 823 cattle (*Bos taurus*) and 188 goats (*Capra aegagrus hircus*) collected in the Eastern Province of Rwanda.

### Reverse Transcription Polymerase Chain Reaction RNA extraction and cDNA synthesis

We extracted the RNA manually using the PureLink Viral RNA/DNA Kit and kit manual. We added 200 μL of OPF sample to a Kit Master Mix (proteinase K, lysis buffer, carrier RNA, and 100% ethanol). We performed a two-step wash using a wash buffer solution, and eluate in 60 μL nuclease-free water was collected and transferred to 1.5 ml tubes. We treated the eluate with DNase using the Turbo DNA-free Kit and kit manual, to remove host genomic DNA. Thereafter, we collected 50 μL of DNA-free eluate and transferred it to 1.5 mL tubes for downstream analyses. We quantified the nucleic material in the collected solutions using the Quantus™ Fluorometer. The SuperScript VILO cDNA Synthesis Kit and kit manual were used for the conversion of RNA extracted manually, and 20 μL from that extracted using the KingFisher Duo machine, to cDNA. We added 10 μL of RNA eluate to a master mix containing 5X VILO reaction mix, 10X SuperScript enzyme mix and nuclease-free water. Complementary DNA synthesis was achieved at 42°C for 60 minutes. Thereafter, we cleaned up the cDNA using the Macherey-Nagel™ Nucleospin Gel and PCR Clean-up Kit and kit manual. Finally, we added 50 μl cDNA to Buffer NE, Buffer NT1, and Buffer NT3 following the methodology detailed in the kit manual.

### RT-PCR

#### Primers

We targeted four serotypes based on the epidemiological status of FMDV pool IV in general and Rwanda in particular (48). We used the primers to detect several FMD serotypes (SAT 1, SAT 2, O and A) both in field samples all purchased from Macrogen (Macrogen Europe, Amsterdam, Netherlands). The RT-PCR primers used in this study are described in the table below.

#### Amplification

The OP samples from the 2017 FMD outbreak were analysed using a multiplex one-step RT-PCR assay. The assay was performed using the TaqMan® Fast Virus 1-Step Kit (Life Technologies) and kit manual. We added 2 μL of eluate extracted in the manual extraction section to a Master Mix containing nuclease-free water, 4X TaqMan buffer, primers and TaqMan probe. The thermal cycler (BioRad, Hercules, California) was programmed as follows; reverse transcription at 50°C for 5 minutes; polymerase activation and DNA denaturation at 95°C for 20 seconds; two-step amplification for 40 cycles with denaturation at 95°C for 3 seconds; annealing and plate read at 60°C for 30 seconds.

We analysed the PCR reaction to observe any positive FMD identification from the OPF samples collected and to compare the results obtained to those from the positive high and low standard controls. Reactions were considered positive if the amplification was detected before 32.0 cycles (50). The blood samples from the 2020 FMD outbreak were analysed using a one-step PCR assay using the QIAGEN One-step RT-PCR Kit (Qiagen GmbH, Hilden, Germany) following the manufacturers’ instructions. Visualization and specification of bands were performed using a 2% agarose gel run at 70 mV for 1 hour.

Before sequencing, we removed the primers and deoxynucleoside by enzymatically adding the ExoSAP-IT PCR Product Cleanup Reagent (0.5μl of Exo and 2μl of SAP) to 10μl of amplicons. The mix was incubated at 37°C for 15 minutes and later 85°C for 15 to activate Exo and SAP respectively, lastly, the product was held at 4°C.

### Real-time reverse transcription loop-mediated isothermal amplification (rRT-LAMP)

Amplification was performed on the Axxin T8–ISO instrument at 65°C for 30 minutes. The assays were performed using the Isothermal Master Mix. Briefly; 6.5 μL cDNA was added to an FMD LAMP primer mix and isothermal master mix.

Amplified products were detected both in real-time and by running an electrophoresis-based gel. The 2% agarose gel bands were read under U.V. light after staining with ethidium bromide. Primers and probes were used according to the manufacturer’s instructions (Tokabio (Pty) Ltd, Johannesburg, South Africa). They are available upon request from the corresponding author.

### Field sample sequencing

We conducted the Whole Genome sequencing of the African buffaloes’ samples without necessarily running a genome-specific PCR amplification. We processed the buffalo samples with the PureLink® Viral RNA/DNA Mini Kit (Invitrogen, Carlsbad, CA) and treated them with the Turbo DNA-free™ Kit (Ambion, TX, USA) to remove any residual host DNA. We then used the Superscript VILO cDNA Synthesis Kit (Invitrogen, Carlsbad, CA) to generate cDNA from the isolated RNA.

NGS libraries were prepared from the samples with the Ion Xpress™ Plus Fragment Library Builder Kit on the AB Library Builder system (Life Technologies). Each sample was uniquely barcoded during library preparation using the Ion Xpress Barcode Adapters 1–16 Kit. We performed template preparation with the Ion PGM Template OT2 200 Kit and the OneTouch 2 instrument, and the samples were sequenced on the Ion PGM next-generation sequencer using the Ion PGM Sequencing 200 Kit. All reagents and instruments were purchased from Thermo Fisher Scientific, Johannesburg, South Africa. Data were analysed with CLC Genomics Workbench Software v.10 (Qiagen Bioinformatics, Redwood City, CA, USA) and the NCBI database with the BLAST tool (https://blast.ncbi.nlm.nih.gov/Blast.cgi).

## List of abbreviations.

FMDV: Foot-and-Mouth Disease Virus
FMD: Foot-and-Mouth Disease
SAT: Southern African Territories
RT-PCR: Reverse Transcription Polymerase Chain Reaction
RT-LAMP: Reverse Transcription Loop-Mediated Isothermal Amplification
CI: Confidence Interval
OP: Oro-pharyngeal
OPF: Oro-pharyngeal fluids
RFU: Relative Fluorescence Units
OIE: Organisation Internationale des Epizooties
NSP: non-structural proteins
ELISA: Enzyme-Linked Immunosorbent Assay
ANP: Akagera National Park
LMICs: Low-and-Middle Income Countries
IM: Intramuscular
EDTA: Ethylenediamine tetra-acetic acid
NCBI: National Centre for Biotechnology Information.

## Consent for publication

Not applicable.

## Competing interests

The authors declare that there are no competing interests.

## Funding

This material is based upon work supported by the United States Agency for International Development, as part of the Feed the Future initiative, under the CGIAR Fund, award number BFS-G-11-00002, and the predecessor fund the Food Security and Crisis Mitigation II grant, award number EEM-G-00-04-00013.

## Authors’ contributions

All authors participated in the manuscript writing. JCU, GOA, GOO, PJL participated in the project designing. JCU, AI and PJL participated in collecting samples. JCU, EU, AI and PJL participated in molecular and serological analyses. PJL and NB participated in sequencing.

## Acknowledgements

The authors thank the African Parks management and staff in Rwanda that granted access to African buffaloes inside the Akagera National Park. Special thanks go to Dr Pete Morkel who helped to tranquillize African buffaloes.

## Author information

1. Centre for Biotechnology and Bioinformatics, University of Nairobi, P.O. Box 30197, Nairobi, Kenya Jean Claude Udahemuka, Gabriel Oluga Aboge & George Ogello Obiero
2. Department of Veterinary Medicine, University of Rwanda, P.O. Box 57, Nyagatare, Rwanda Jean Claude Udahemuka & Evodie Uwibambe
3. Department of Public Health, Pharmacology and Toxicology, Faculty of Veterinary Medicine, University of Nairobi, P.O. Box 29053, Nairobi, Kenya Gabriel Oluga Aboge
4. Rwanda Agriculture and Animal Resources Board, P.O. Box 5016 Huye, Rwanda. Ingabire Angelique
5. TokaBio (Pty), Ltd, Pretoria, South Africa Phiyani Justice Lebea & Natasha Beeton

## Notes

### Competing Interest Statement

The authors have declared no competing interest.

